# SARS-CoV-2 nucleocapsid protein undergoes liquid-liquid phase separation stimulated by RNA and partitions into phases of human ribonucleoproteins

**DOI:** 10.1101/2020.06.09.141101

**Authors:** Theodora Myrto Perdikari, Anastasia C. Murthy, Veronica H. Ryan, Scott Watters, Mandar T. Naik, Nicolas L. Fawzi

**Affiliations:** Center for Biomedical Engineering, Brown University, Providence, RI, USA; Molecular Biology, Cell Biology & Biochemistry Graduate Program, Brown University, Providence, RI, USA; Neuroscience Graduate Program, Brown University, Providence, RI, USA; Department of Molecular Pharmacology, Physiology, and Biotechnology, Brown University, Providence, RI, USA; Robert J. and Nancy D. Carney Institute for Brain Science, Brown University, Providence, RI, USA

## Abstract

Tightly packed complexes of nucleocapsid protein and genomic RNA form the core of viruses and may assemble within viral factories, dynamic compartments formed within the host cells. Here, we examine the possibility that the multivalent RNA-binding nucleocapsid protein (N) from the severe acute respiratory syndrome coronavirus (SARS-CoV-2) compacts RNA via protein-RNA liquid-liquid phase separation (LLPS) and that N interactions with host RNA-binding proteins are mediated by phase separation. To this end, we created a construct expressing recombinant N fused to a N-terminal maltose binding protein tag which helps keep the oligomeric N soluble for purification. Using *in vitro* phase separation assays, we find that N is assembly-prone and phase separates avidly. Phase separation is modulated by addition of RNA and changes in pH and is disfavored at high concentrations of salt. Furthermore, N enters into *in vitro* phase separated condensates of full-length human hnRNPs (TDP-43, FUS, and hnRNPA2) and their low complexity domains (LCs). However, N partitioning into the LC of FUS, but not TDP-43 or hnRNPA2, requires cleavage of the solubilizing MBP fusion. Hence, LLPS may be an essential mechanism used for SARS-CoV-2 and other RNA viral genome packing and host protein co-opting, functions necessary for viral replication and hence infectivity.

## Introduction

The spread of the highly infectious severe acute respiratory syndrome coronavirus 2 (SARS-CoV-2) is responsible for the ongoing global pandemic of <u>Co</u>rona<u>v</u>irus <u>Di</u>sease 20<u>19</u> (COVID-19)^1^. The novel SARS-CoV-2 is an enveloped, nonsegmented, positive-sense, single stranded ~30 kb RNA virus of the family *Coronaviridae*^2^. This family includes the related SARS-CoV^3^ and Middle East respiratory syndrome (MERS)^4^ coronaviruses, which have both caused previous outbreaks of pneumonia. Like all coronaviruses, SARS-CoV-2 forms a virion including its genomic RNA (gRNA) packaged in a particle comprised of four structural proteins – the crownlike spike (S) glycoprotein that binds to human ACE2 receptor to mediate the entry of the virus in the host cell^5,6^, the membrane (M) protein that facilitates viral assembly in the endoplasmic reticulum, the ion channel envelope (E) protein, and the nucleocapsid protein (N) that assembles with viral RNA to form a helical ribonucleoprotein (RNP) complex called the nucleocapsid^7,8^. Though many current therapeutic efforts have focused on disrupting viral attachment to host cells^9^ and preventing viral protease function^10^, the molecular mechanisms that underlie the assembly of the SARS-CoV-2 nucleocapsid through the binding of nucleoprotein to RNA are poorly understood and therefore have remained an uninvestigated target to inhibit viral replication.

Nucleocapsid protein of SARS-CoV-2 is a 46 kDa multivalent RNA-binding protein which is predicted to have 40% of its primary sequence remaining intrinsically disordered in addition to the two known folded domains. N has a folded N-terminal domain that participates in RNA-binding domain (NTD) preceded by a 44-amino acid N-terminal disordered region (N_IDR_) and followed by a serine/arginine-rich 73-amino acid linker (LKR_IDR_) (**Figure 1A**). The flexible linker is adjacent to the folded dimerization domain (CTD) followed by the 52-amino C-terminal disordered tail region (C_IDR_) rich in lysine and glutamine (**Figure 1B**). Previous studies of SARS-CoV-1 N (91% sequence identity) have shown that both the NTD, the CTD and the disordered regions can bind RNA cooperatively to promote RNP packaging^11^ and chaperoning^12^. The structural details of these protein-RNA interactions are beginning to come into focus. A recent solution NMR structure of SARS-CoV-2 NTD-RNA complex suggested a right hand-like fold and highlights the role of arginine motifs and electrostatic interactions^13^. Furthermore, superimposition of the unbound state with previously solved coronavirus nucleoproteins bound to RNA revealed some potential protein-RNA recognition interactions involving Arg89, Tyr110 and Tyr112 in nitrogenous base binding^14^. The role of NTD in viral genome packaging has been investigated in HCoV-OC43 N-NTD^15^ and mouse hepatitis virus (MHV) N-NTD which organizes gRNA via specific interactions with a packaging signal (PS) located 20.3 kb from the 5’ end of gRNA^16^. Importantly, the higher-order assembly of coronavirus N proteins has also begun to be examined. The CTD dimerization domain of SARS-CoV has been suggested to organize into an octamer stabilized via electrostatic interactions enhanced by phosphorylation^17^ to promote superhelical packaging of viral RNA^18^.

**Figure 1:**
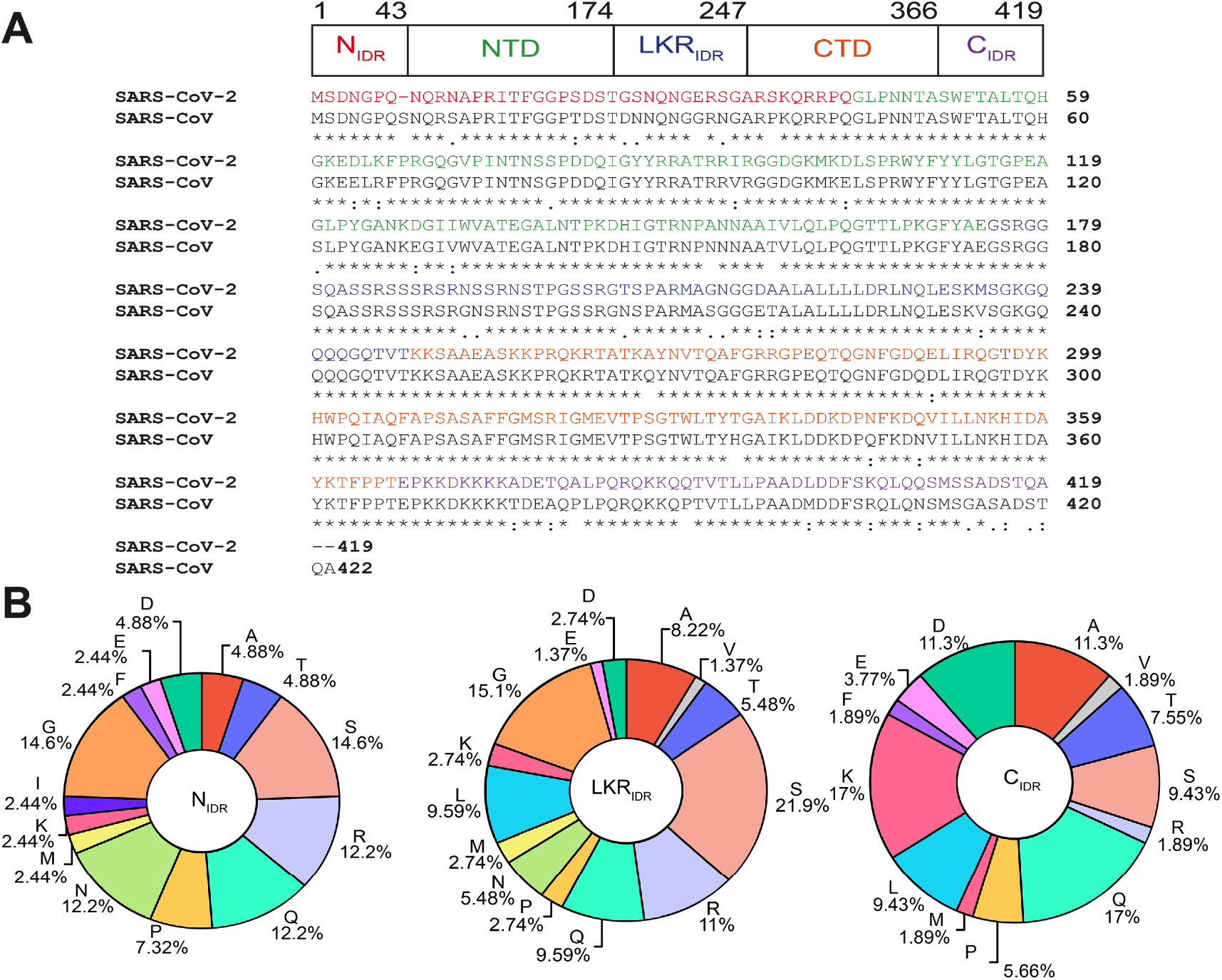
Domain structure, sequence comparison, and sequence composition of SARS-CoV-2 nucleocapsid protein. A) SARS-CoV-2 N contains three putatively disordered regions, a globular N-terminal and a globular C-terminal oligomerization domain. Sequence alignment of N from SARS-CoV-2 and SARS-CoV showing 91% sequence identity. B) Sequence composition of the intrinsically disordered regions of SARS-CoV-2 N.

In recent years, liquid-liquid phase separation (LLPS) has emerged as a common cellular process to organize biological material into compartments. Many of these biomolecular condensates assembled by LLPS are multicomponent phases composed of multivalent RNA-binding proteins containing intrinsically disordered regions (IDRs) and RNA^19^. Biomolecular condensates are thought to be formed by weak, multivalent interactions to sequester and concentrate proteins involved in RNA processing, stress response and gene silencing^20^. In eukaryotes, histone proteins are known to promote the compaction of chromatin into condensates^21^. Bacterial nucleoprotein complexes also have been shown to organize genomic DNA via phase separation^22^, suggesting that LLPS may serve to organize genome packaging across the domains of life. Recent evidence suggests that similar higher order genome organization is also present in viruses. For example, the measles virus nucleoprotein (MeV N) assembles with genomic RNA into a rigid helical capsid^23^ which also undergoes LLPS in the presence of the phosphoprotein (P)^24^. Like the eukaryotic heterochromatin protein 1 (HP1) which bridges chromatin regions and can undergo LLPS^25^, SARS-CoV-2 N is oligomeric with multiple binding sites for the genomic nucleic acid separated by disordered linkers. Hence, it is important to understand if SARS-CoV-2 N could also use phase separation to organize its genome.

Furthermore, viruses have long been known to hijack the host cell environment to facilitate gRNA transport from the site of viral genome replication to the site of viral assembly and maximize the replication efficiency by disrupting the organization of cellular organelles^26,27^. A recent study on the network of protein-protein contacts formed by SARS-CoV-2 structural proteins shows that N interacts with several human ribonucleoproteins known to be involved in the formation of phase-separated protein-RNA granules^28^. Several of these proteins such as G3BP1/2 are composed of disordered regions with sequence characteristics that are known to contribute to stress granule formation via LLPS^29,30^. Stress granules (SGs) are membraneless organelles that store translationally silent mRNA when the cell is exposed to stress to regulate mRNA metabolism^31^. Numerous studies have shown viral invasion can interfere with SG formation^32^ via inhibition of post translational modifications^33^, exclusion of SG components such as TIA-1 and G3BP^34,35^, and formation of stable viral RNP complexes with SG vital proteins^36^. Hence, it would be important to understand if SARS-CoV-2 N can enter phase-separated assemblies formed by other ubiquitous, well-characterized SG human proteins. Here, we examine the phase separation of N *in vitro* as a function of solution condition and RNA concentration. Furthermore, we test the partitioning of N into phase separated droplets formed by intact human ribonucleoproteins or their disordered domains.

## Results

### N forms higher order oligomers

The SARS-CoV nucleocapsid protein contains multiple regions implicated in self-interactions^17,37–39^, so we first sought to determine if the SARS-CoV-2 nucleocapsid protein (N) was capable of higher order assembly. We purified recombinant full-length SARS-CoV-2 N with a TEV protease cleavable N-terminal maltose binding protein (MBP) tag to enhance solubility, due to previous studies showing that the N RNA complex was largely insoluble^11^. Even with the solubility tag, some of the N protein deposited into inclusion bodies during bacterial expression (data not shown). However, purification of MBP-N from the soluble fraction was efficient at high salt concentration via standard immobilized metal affinity chromatography. The chromatogram from subsequent gel filtration chromatography of both MBP-tagged and cleaved (tag removed) N have major absorbance peaks eluting much earlier than would be expected of their monomeric species (90 kDa, expected at about 230 mL and 46 kDa expected at about 240 mL, respectively) (**Figure 2A**). When analyzed by SDS-PAGE, the fractions containing these peaks correspond to the MBP-N and cleaved N respectively with some minor degradation products but no species of greater molecular weight (**Figure 2B, C**). These data imply that N is capable of stable self-assembly.

**Figure 2:**
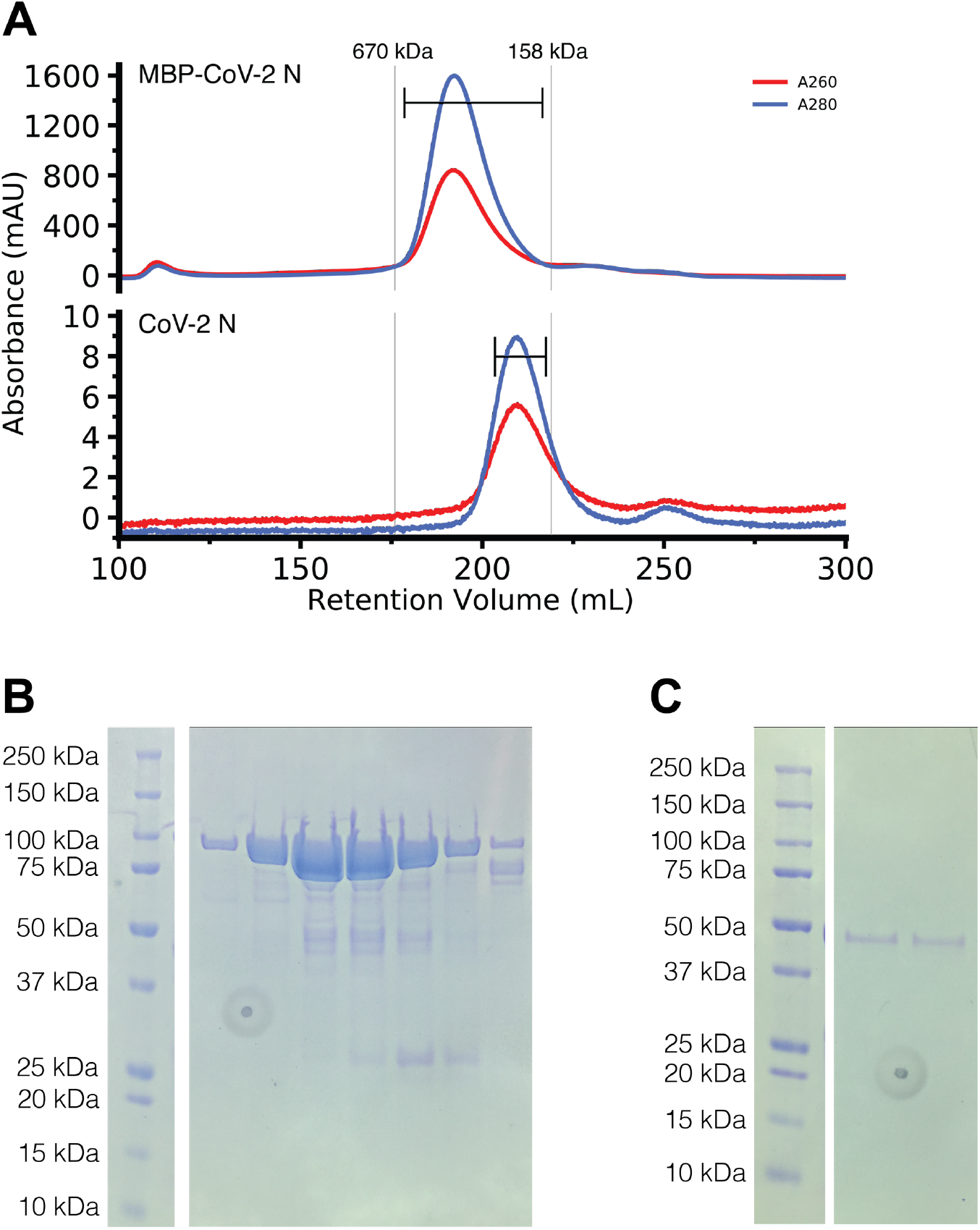
MBP-CoV-2 N and cleaved CoV-2 N elute larger than their predicted molecular weights. A) Gel filtration chromatogram of ~7ml of ~500 μM MBP tagged CoV-2 N (top) and ~300μl ~50 μM CoV-2 N (previously cleaved from MBP) (bottom). Vertical lines represent peak elution volumes of gel filtration protein calibration standards. Brackets represent range of gels lanes. Predicted molecular weight of the tagged and untagged N are 90 kDa and 46 kDa respectively, much smaller than their corresponding calibrated peak elution volumes, consistent with oligomerization. B) SDS-PAGE showing the peak fractions of the MBP-CoV-2 N chromatogram showing expected 90 kDa MW. C) SDS-PAGE showing the peak fractions of the cleaved CoV-2 N chromatogram showing expected 46 kDa MW. Circular feature in SDS-PAGE gels is a divet in plastic on which pictures were taken.

### N undergoes LLPS *in vitro*

The SARS-CoV nucleocapsid protein associates with viral genomic RNA to form a ribonucleoparticle and is enriched in serine and arginine residues^40,41^, a characteristic of some proteins that can undergo LLPS. We hypothesized that N is able to form liquidlike compartments to sequester RNA. To test if N is able to phase separate, we prepared samples containing MBP-tagged full-length N at a variety of conditions, added TEV, and measured the resulting turbidity of the solution coupled with microscopy as we have done for previous studies of LLPS-prone proteins^42^. First, we tested whether N could undergo LLPS in a variety of pH conditions in the presence and absence of torula yeast RNA extract (desalted to remove ions and small RNA pieces) (**Figure 3**). Like in our previous work examining the impact of RNA binding on phase separation of FUS which binds many RNA sequences and structures^43^, we used RNA extract as N has been shown to bind with little specificity to nucleic acids including ssRNA, ssDNA and dsDNA^11,38^. At pH 7.4, mixtures containing 50 μM MBP-N with 0.3 mg/mL RNA in the presence of TEV protease displays initial increased turbidity over time (**Figure 3A**) followed by a decrease, characteristic for the formation of turbid droplet assemblies triggered by TEV cleavage which then fuse and settle^44,45^. This increase in turbidity is coupled with the appearance of small, spherical droplets visible by microscopy (**Figure 3B**). Interestingly, at lower pH conditions the turbidity of the reaction increases and persists over time as has been observed for protein aggregates or gels^46^ (**Figure 3A**). At pH 5.5, the droplets still appear spherical; however, the persistent turbidity suggests that the droplets may be less fluid (more “gel-like”) in these conditions. Consistent with this hypothesis, at lower pH conditions the assemblies no longer remain spherical and resemble droplets that were unable to complete fusion (**SI Figure 1A**). Further, addition of MgCl_2_ or CaCl_2_ does not substantially alter LLPS of N (**SI Figure 2**), demonstrating that presence of divalent metal ions does not affect LLPS. Together, these data suggest that SARS-CoV2 N is able to undergo LLPS at physiological buffer conditions.

**Figure 3:**
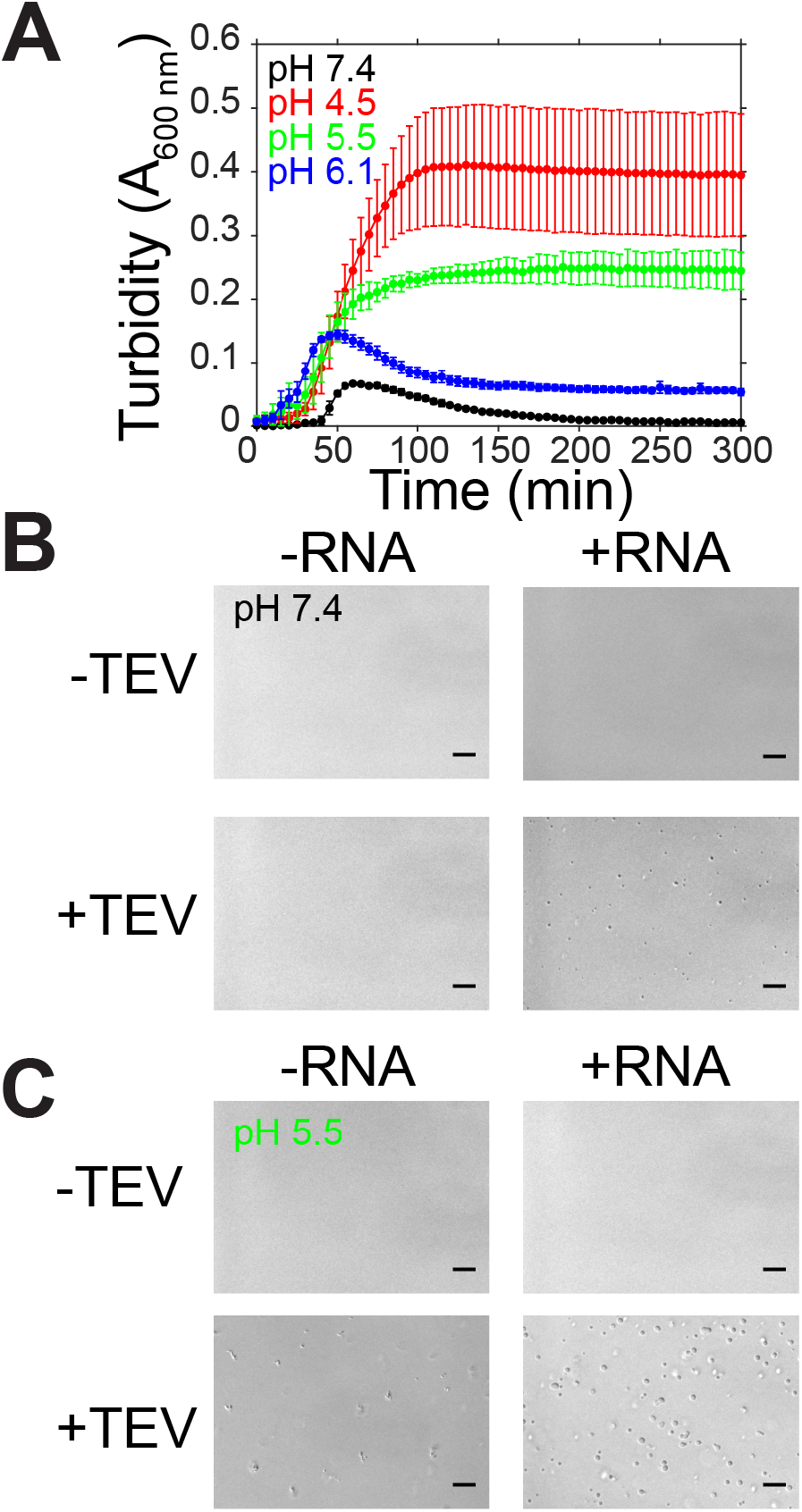
SARS-CoV-2 nucleocapsid protein undergoes LLPS at physiological conditions. A) Phase separation over time as monitored by turbidity of 50 μM MBP-N after addition of TEV protease in varying pH conditions. B-C) DIC micrographs of 50 μM MBP-N in 50 mM Tris 183 mM NaCl pH 7.4 or 20 mM MES 183 mM NaCl pH 5.5 with and without TEV protease (to cleave MBP from N) or 0.3 mg/mL desalted total torula yeast RNA. Scale bars represent 50 μm.

### N LLPS is modulated by RNA concentration and ionic strength

Because RNA is involved in the assembly of the viral RNP, we sought to determine if RNA concentration and interactions were important for droplet assembly. We conducted turbidity and microscopy experiments with fixed protein concentration and varying protein:RNA mass ratios in low salt conditions (**Figure 4A,C**). In low salt conditions in the absence of RNA, there is an increase in turbidity along the formation of small, spherical droplets, suggesting that N is able to undergo LLPS even in the absence of RNA (**Figure 4A,C**). At conditions of 1:0.25 MBP-N:RNA, the turbidity of the solution is enhanced compared to the no RNA control (**Figure 4A**). Interestingly, at higher RNA concentrations turbidity and droplet formation are diminished (**Figure 4A,C**), a characteristic of reentrant phase separation behavior^47^. To test if the electrostatic interactions between N and RNA are important for phase separation, we measured turbidity accompanied by microscopy of solutions containing varying sodium chloride concentrations (**Figure 4B,D**). We found that at higher salt concentrations (300 mM and 1M), both turbidity and droplet formation was reduced (**Figure 4B**). Together, these data show protein-RNA interactions stimulate N phase separation and suggest that screening out the electrostatic interactions between N and RNA reduces LLPS.

**Figure 4:**
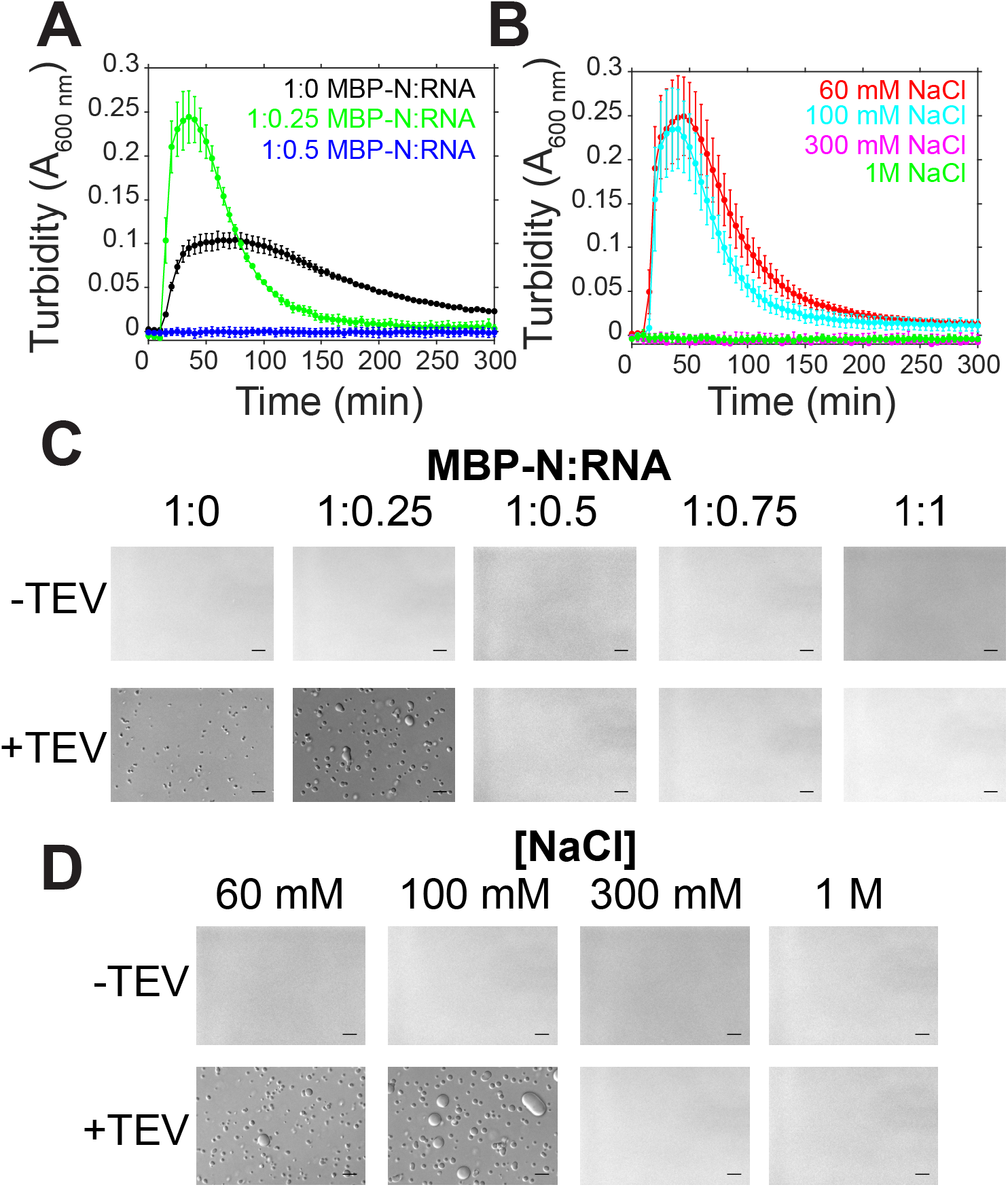
SARS-CoV-2 N LLPS is modulated by salt and RNA. A-B) Phase separation over time as monitored by turbidity of 50 μM MBP-N in 50 mM Tris pH 7.4 after addition of TEV protease with varying torula yeast RNA (at 100 mM sodium chloride) or varying sodium chloride concentrations. C-D) DIC micrographs of 50 μM MBP-N in 50 mM Tris NaCl pH 7.4 with varying torula yeast RNA or sodium chloride concentrations with and without TEV protease (to cleave MBP from N). Scale bars represent 50 μm.

### N partitions into hnRNPA2, TDP-43, and FUS droplets

SARS-CoV-2 N interacts with stress granule proteins^28^ and the interaction between one stress granule protein, hnRNPA1, and SARS-CoV-1 N has been examined with biochemical detail^37^. As we have previously shown co-partitioning of many granule-associated heterogenous nuclear ribonucleoproteins (hnRNPs) into liquid phases including intermixing of FUS, hnRNPA2, and TDP-43^45,48^, we decided to test if full length N could partition into liquid phases formed by hnRNPA2, FUS, and TDP-43. We used conditions where N does not phase separate on its own (i.e. ~5 nM concentration with no RNA) to be sure that N was the “client” and the hnRNP was the “scaffold” protein, following common terminology to classify the molecules as drivers (“scaffold”) or partitioning members (“clients”) of phase separation, respectively^49,50^. We found that N partitions into hnRNPA2 LC and TDP-43 CTD droplets even when attached to the maltose binding protein (MBP) solubility tag (**Figure 5A-B**). In contrast, MBP-N did not partition into FUS LC (**Figure 5C**), suggesting a weaker interaction with FUS LC than with TDP-43 CTD or hnRNPA2 LC. However, N did partition into FUS LC droplets when cleaved from MBP (**Figure 5C**). We further tested whether N could partition into droplets formed by the full-length hnRNPs. We found that N was able to partition into full-length hnRNPA2, FUS, and TDP-43 droplets (**Figure 5D-F**). As the hnRNPs are only able to undergo LLPS after cleavage of the MBP solubility tag, we could not observe partitioning of MBP-N and hnRNPs. These results indicate that N can interact with and partition into liquid phases formed by many human RNA-binding proteins, suggesting the presence of weak protein-protein interactions.

**Figure 5:**
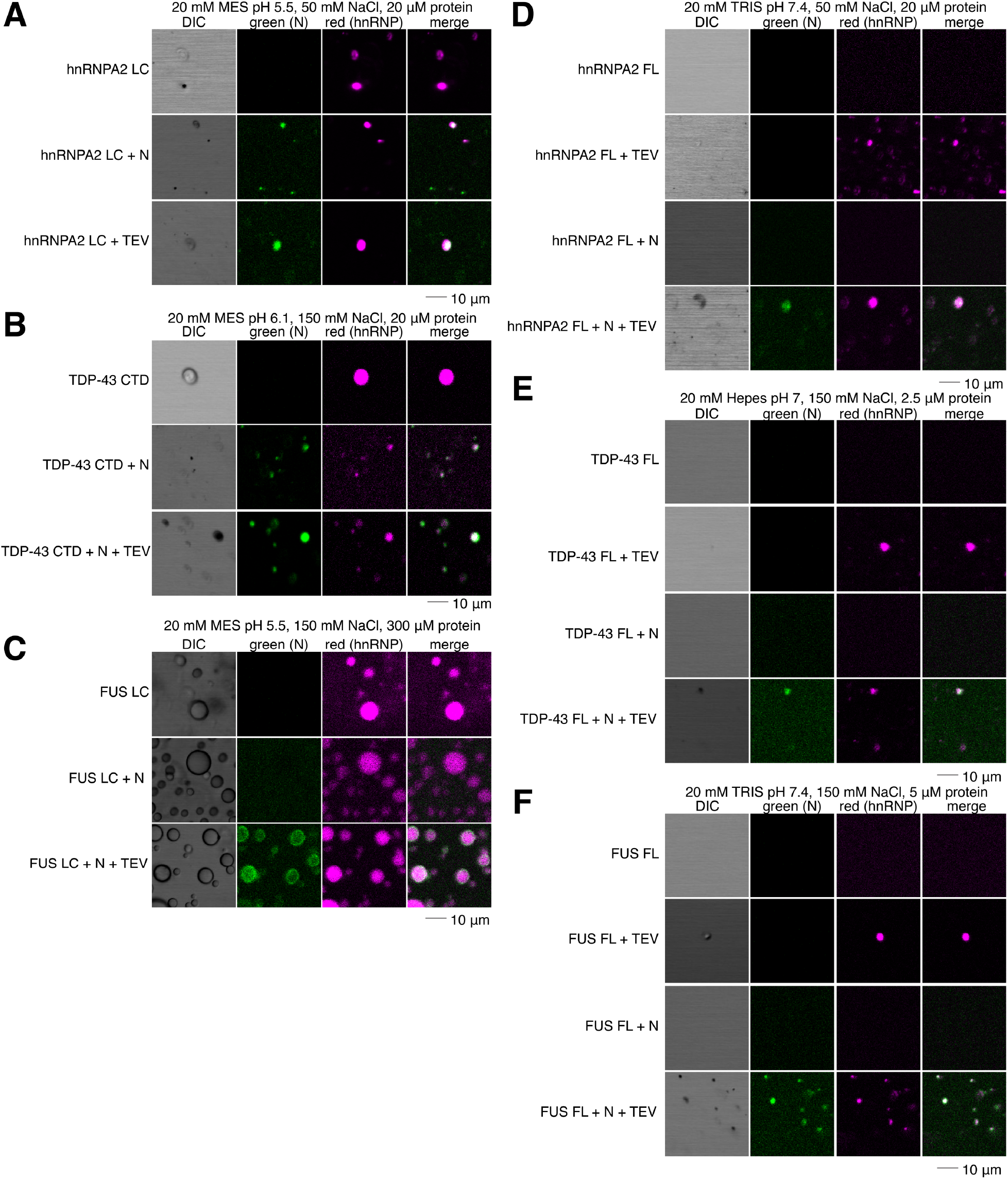
SARS-CoV-2 N partitions into liquid phases of hnRNPs. A) N partitions into hnRNPA2 LC droplets even with the MBP tag attached. B) N partitions into TDP-43 CTD droplets even with the MBP tag attached. C) N partitions into FUS LC droplets only after cleavage of the MBP tag. D-F) N partitions into hnRNPA2 FL (D), TDP-43 FL (E), FUS FL (F) droplets.

## Discussion

Virion assembly requires the formation of dense protein-nucleic acid compartments that sequester host cell proteins as a means of protection from the host immune system and concentrate viral components to increase the efficiency of replication^26^. Previous studies on herpes virus have reported the existence of perinuclear and cytoplasmic puncta in infected cells called prereplicative sites^51,52^. Apart from SARS CoV-1 and SARS-CoV-2, a vast number of viruses like paramyxoviruses^53^, flaviviruses^54^, and mononegaviruses such as rabies^55,56^, influenza A virus^57,58^ and Lassa virus^59^ are also known to hijack their host cellular machinery via their highly disordered nucleoprotein with some of them resulting to the formation of cytoplasmic puncta such as the Negri bodies of *Mononegavirales*^56^. Most recently, the co-phase separation of full-length nucleoprotein (N) and phosphoprotein (P) of measles virus (MeV) was shown to require both the folded and the disordered domain of N, highlighting the importance of LLPS during MeV replication^24^. Understanding the mechanisms that underlie LLPS of SARS-CoV-2 N is essential to identify key potentially targetable steps in the viral replication cycle. To unravel the molecular details of this phenomenon, we tested the ability of N protein to undergo phase separation in the presence of RNA and other human ribonucleoproteins that are located in membraneless organelles of the eukaryotic cytoplasm and nucleoplasm^60^.

Here, we found that N of SARS-CoV-2 is able to phase separate and its phase behavior is tuned by pH, salt, and RNA concentration. At physiological conditions (pH = 7.4, NaCl = 183 mM), we observed the formation of *in vitro* N droplets after addition of TEV protease and 0.3 mg/ml desalted total torula yeast RNA. Remarkably, at lower pH values, the turbidity of the RNA-protein mixtures increased monotonically, and the droplets persisted, albeit the sphericity and the fluidity were diminished (**SI Figure 1**). After determining conditions where N phase separates, we tested the dependence of LLPS on RNA and salt concentration. Surprisingly, we found in the absence of RNA at 100 mM NaCl, N formed small spherical droplets. At 1:0.25 protein:RNA ratio, the size and the sphericity of the droplets further increased. However, higher RNA ratios eliminated phase separation, suggesting that N of SARS-CoV-2 exhibits RNA-mediated phase-behavior typical of many liquid-like transcriptional machineries with reentrant phase transition^47^. Lastly, we tested the effect of ionic strength on LLPS of N and we showed that at higher salt concentrations (300 mM, 1 M), N no longer phase separates as supported by our microscopy and turbidity data. It is unclear which domains of N are essential for phase separation, although previous studies on N of SARS-CoV-1 have shown that the NTD^18^, the CTD^61^, and the disordered linker can bind RNA cooperatively with a degree of specificity being attributed to the positive electrostatic surface of these regions. Thus, here we demonstrate that the nucleocapsid protein from a virus of the *Coronaviridae* family undergoes phase separation with RNA *in vitro*.

Recent proteomic studies have constructed a putative SARS-CoV-2 protein interaction map where many RNA processing factors and stress granule regulation factors like G3BP1/2^29^ have been delineated as crucial nodes of the N interactome^28^. In addition, early reports on molecules interacting with N of SARS-CoV-1 highlighted high binding affinity to granule-associated proteins – hnRNPA1^37^, the stress-granule and phase-separating protein^62^; nucleophosmin (nucleolar phosphoprotein B23)^63^, the primary component of the phase separated liquid granular component of the nucleolus^64^, and cyclophilinA^65^, the hnRNP chaperone^66^. During the steps of genomic replication in the infected cells, the local concentration of cellular organelles is altered dramatically by the shuttling of the replication complexes in the cytoplasm and the anchorage of the structural proteins to the cellular membranes^7^. We hypothesized that N facilitates SARS-CoV-2 replication by recruiting stress granule components present in the host cellular environment. Here, we identified the interaction between full-length N of SARS-CoV-2 and human ribonucleoproteins like FUS, hnRNPA2 and TDP-43 full-length and their respective low complexity domains. It is possible that abundant host cytoplasmic proteins, like these hnRNPs, serve as scaffolds to promote the formation of multicomponent dense N-RNA phases to enable or accelerate viral replication. Given that we have shown here the potential role of phase separation in stabilizing nucleocapsid formation and the ability of viral nucleocapsid proteins to enter phase separated assemblies formed by host cell RNA-binding proteins, it will be important to investigate if therapies targeting viral or host condensates could disrupt cycles of SARS-CoV-2 replication.

## Acknowledgements

We thank Gerwald Jogl and Walter Atwood for helpful input and Geoff Williams and Christoph Schorl for technical assistance. We thank Abigail Janke and Alexander Conicella for making available stocks of their FUS and TDP-43 full-length protein, respectively. Research was supported by a COVID-19 Research Seed Award from Brown University (to M.T.N., N.L.F, Gerwald Jogl, and Walter Atwood), the Division of Biology and Medicine at Brown University, the National Institute of General Medical Sciences R01GM118530 (to N.L.F.), the National Institute of Neurological Diseases and Stroke and the National Institute on Aging R01NS116176 (to N.L.F.). A.C.M. was supported by an NSF graduate fellowship (1644760). V.H.R. was supported by the National Institutes of Health F31NS110301.

## Author contributions

T.M.P performed bioinformatics analysis and wrote the introduction and discussion. A.C.M. performed phase separation turbidity assays and microscopy of N. V.H.R performed N partitioning assays and turbidity screening of divalent metal salts. S.W. designed the expression construct and purified the protein. N.L.F. and M.T.N. contributed to research design and funding acquisition. T.M.P led the writing of the manuscript with text, figures and comments provided by all authors (V.H.R, A.C.M, S.W, M.T.N. and N.L.F).

## Methods

### Constructs

- MBP-SARS-CoV-2 N full-length, soluble histag purification
- hnRNPA2 LC, insoluble histag purification (Addgene: 98657)
- TDP-43 CTD, insoluble histag purification (Addgene: 98670)
- FUS LC, insoluble anion purification (Addgene: 98656)
- MBP-hnRNPA2 FL, soluble histag purification (Addgene: 139109)
- MBP-TDP-43 FL, soluble histag purification (Addgene: 104480)
- MBP-FUS FL, soluble histag purification (Addgene: 98651)

### MBP-N full-length expression and purification

MBP-tagged (pTHMT) full-length SARS-CoV2 nucleocapsid protein was expressed in *Escherichia coli* BL21 Star (DE3) cells (Life Technologies). Bacterial cultures were grown to an optical density of 0.7 to 0.9 before induction with 1 mM isopropyl-β-D-1-thiogalactopyranoside (IPTG) for 4 hrs at 37°C. Cell pellets were harvested by centrifugation and stored at −80°C. Cell pellets were resuspended in 20 mM Tris 1M NaCl 10 mM imidazole pH 8.0 with one EDTA-free protease inhibitor tablet (Roche) and lysed using an Emulsiflex C3 (Avestin). The lysate was cleared by centrifugation at 20,000 rpm for 50 min at 4°C, filtered using a 0.2 μm syringe filter, and loaded onto a HisTrap HP 5 mL column. The protein was eluted with a gradient from 10 to 300 mM imidazole in 20 mM Tris 1.0 M NaCl pH 8.0. Fractions containing MBP-N full-length were loaded onto a HiLoad 26/600 Superdex 200 pg column equilibrated in 20 mM Tris 1.0 M NaCl pH 8.0. Fractions with high purity were identified by SDS-PAGE and concentrated using a centrifugation filter with a 10 kDa cutoff (Amicon, Millipore).

### N full-length cleavage from MBP, MBP-tag removal, and N gel filtration

MBP-N was incubated at ~600 μM in 20 mM Tris 1.0 M NaCl pH 8.0 with 0.03 mg/mL in-house TEV protease overnight. The protein was then buffer exchanged into 20 mM Tris 1.0 M NaCl pH 8.0 10 mM imidazole using a centrifugation filter with a 10 kDa cutoff (Amicon, Millipore). MBP (and his-TEV) was then removed (“subtracted”) using a HisTrap HP 5 mL column and flow through fractions containing cleaved N full-length were loaded onto a HiLoad 26/600 Superdex 200 pg column equilibrated in 20 mM Tris 1.0 M NaCl pH 8.0. Fractions from gel filtration were analyzed by SDS-PAGE.

### hnRNP purification

hnRNPA2 LC^45^, MBP-hnRNPA2 FL^48^, TDP-43 CTD^67^, MBP-TDP-43 FL^68^, FUS LC^42^, and MBP-FUS FL^44^ were purified as described.

### AlexaFluor labeling

Proteins were labeled with NHS-ester AlexaFluor dyes by diluting protein stocks into 20 mM HEPES pH 8.3 1 M NaCl (for N, hnRNPA2 FL, TDP-43 FL, and FUS FL) or 20 mM HEPES pH 8 with 8 M urea (FUS LC, hnRNPA2 LC, TDP-43 CTD). AlexaFluor dissolved in DMSO was added at less than 10% total reaction volume. Reactions were incubated for an hour and unreacted AlexaFluor was removed by desalting with 1 mL Zeba spin desalting columns equilibrated in the appropriate buffer for protein solubility. Labeled proteins were then concentrated and buffer exchanged into appropriate storage buffers and flash frozen.

### Turbidity measurements

Turbidity was used to evaluate phase separation of 50 μM MBP-N full-length in the presence of 0.01 mg/mL in-house TEV protease (~0.3 mg/mL in 50 mM Tris 1 mM EDTA 5 mM DTT pH 7.5 50% glycerol 0.1% Triton-X-100) in the appropriate conditions. To test the effect of pH on LLPS, the experiment was conducted in 50 mM Tris 183 mM NaCl pH 7.4, 20 mM MES 183 mM pH 6.1, 20 mM MES 183 mM pH 5.5, 20 mM MES 183 mM pH 4.9, 20 mM MES 183 mM pH 4.5 with 0.3 mg/mL desalted (into the appropriate buffer using a Zeba 0.5 mL spin column) torula yeast RNA extract in the appropriate buffer conditions. To test the effect of different salt concentrations on LLPS, the experiments were conducted in 50 mM Tris pH 7.4 with 60 mM, 100 mM, 300 mM or 1000 mM NaCl with 1.1 mg/mL desalted torula yeast RNA extract. To test the effect of RNA on LLPS, the experiments were conducted in 50 mM Tris 100 mM NaCl pH 7.4 with 0, 1.1 or 2.3 mg/mL desalted torula yeast RNA extract. Turbidity experiments were performed in a 96-well clear plate (Costar) with 70 μL samples sealed with optical adhesive film to prevent evaporation (MicroAmp, ThermoFisher). The absorbance at 600 nm was monitored over time using a Cytation 5 Cell Imaging Multi-Mode Reader (BioTek) at 5 min time intervals for up to 12 hr with mixing. To subtract background noise, the turbidity of a no TEV control (i.e. replaced with TEV storage buffer) for each condition was subtracted from the turbidity of the experimental conditions. Experiments were conducted in triplicate and averaged.

### DIC microscopy

For 50 μM MBP-N full-length, the samples were incubated with 0.03 mg/mL in-house TEV protease for 20 min before visualization. Samples were spotted onto a glass coverslip and droplet formation was evaluated by imaging with differential interference contrast on an Axiovert 200M microscopy (Zeiss).

### hnRNP mixing microscopy

Each protein was prepared for microscopy based on our previously established methods for that protein^42,44,45,48,67,68^. Briefly: hnRNPA2 was diluted from 8 M urea into the appropriate buffer to a final concentration of 150 mM urea and appropriate protein concentration; hnRNPA2 FL with a C-terminal MBP tag was diluted from 1 M NaCl to 50 mM NaCl and appropriate protein concentration; TDP-43 CTD was desalted into MES pH 6.1 using a 0.5 mL Zeba spin desalting column and diluted to the appropriate protein concentration; TDP-43 FL with a C-terminal MBP tag was diluted from storage buffer into appropriate buffer at indicated concentration; FUS LC was was diluted from 20 mM CAPS to appropriate concentration in indicated buffer; and FUS FL with an N-terminal MBP tag was diluted from 1 M NaCl to a final NaCl concentration of 150 mM and indicated protein concentration. 1 μL of 0.3 mg/mL TEV was added as appropriate, if no TEV was needed for the sample, TEV storage buffer was added instead. Buffer conditions for each protein are listed below:

- hnRNPA2 LC: 20 μM hnRNPA2 LC, 20 mM MES pH 5.5, 50 mM NaCl, 150 mM urea (residual), ~5 nM AlexaFluor labeled protein (each, if both N and hnRNP are present)
- hnRNPA2 FL: 20 μm hnRNPA2 FL, 20 mM TRIS pH 7.4, 50 mM NaCl, ~5 nM AlexaFluor labeled protein (each, if both N and hnRNP are present)
- TDP-43 CTD: 20 μM TDP-43 CTD, 20 mM MES pH 6.1, 150 mM NaCl, ~5 nM AlexaFluor labeled protein (each, if both N and hnRNP are present)
- TDP-43 FL: 2.5 μM TDP-43 FL, 20 mM HEPES pH 7, 150 mM NaCl, 1 mM DTT, ~5 nM AlexaFluor labeled protein (each, if both N and hnRNP are present)
- FUS LC: 300 μM FUS LC, 20 mM MES pH 5.5, 150 mM NaCl, ~5 nM AlexaFluor labeled protein (each, if both N and hnRNP are present)
- FUS FL: 5 μM FUS FL, 20 mM TRIS pH 7.4, 150 mM NaCl, ~5 nM AlexaFluor labeled protein (each, if both N and hnRNP are present)

Fluorescence confocal microscopy images were taken on an LSM 880 (Zeiss). AlexaFluor-tagged proteins were doped in at 0.2 μL (~5 nM final concentration) to prevent oversaturation of the detector. Snapshots were taken of the red, green, and brightfield channels and merged using ImageJ (NIH).

---

**SI Figure 1:**
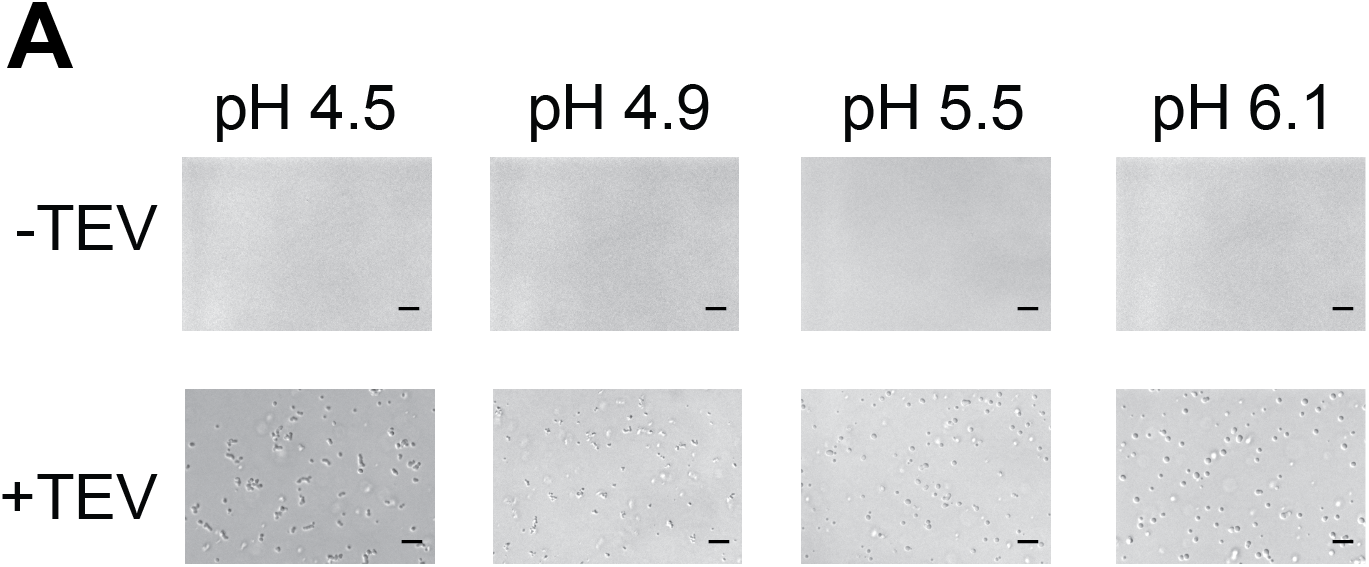
Low pH conditions induce aggregation of MBP-NP. A) DIC micrographs of 50 μM MBP-N in varying pH conditions. At lower pH conditions, droplets appear to be non-spherical, consistent with less fluid behavior. Scale bars represent 50 μm.

**SI Figure 2:**
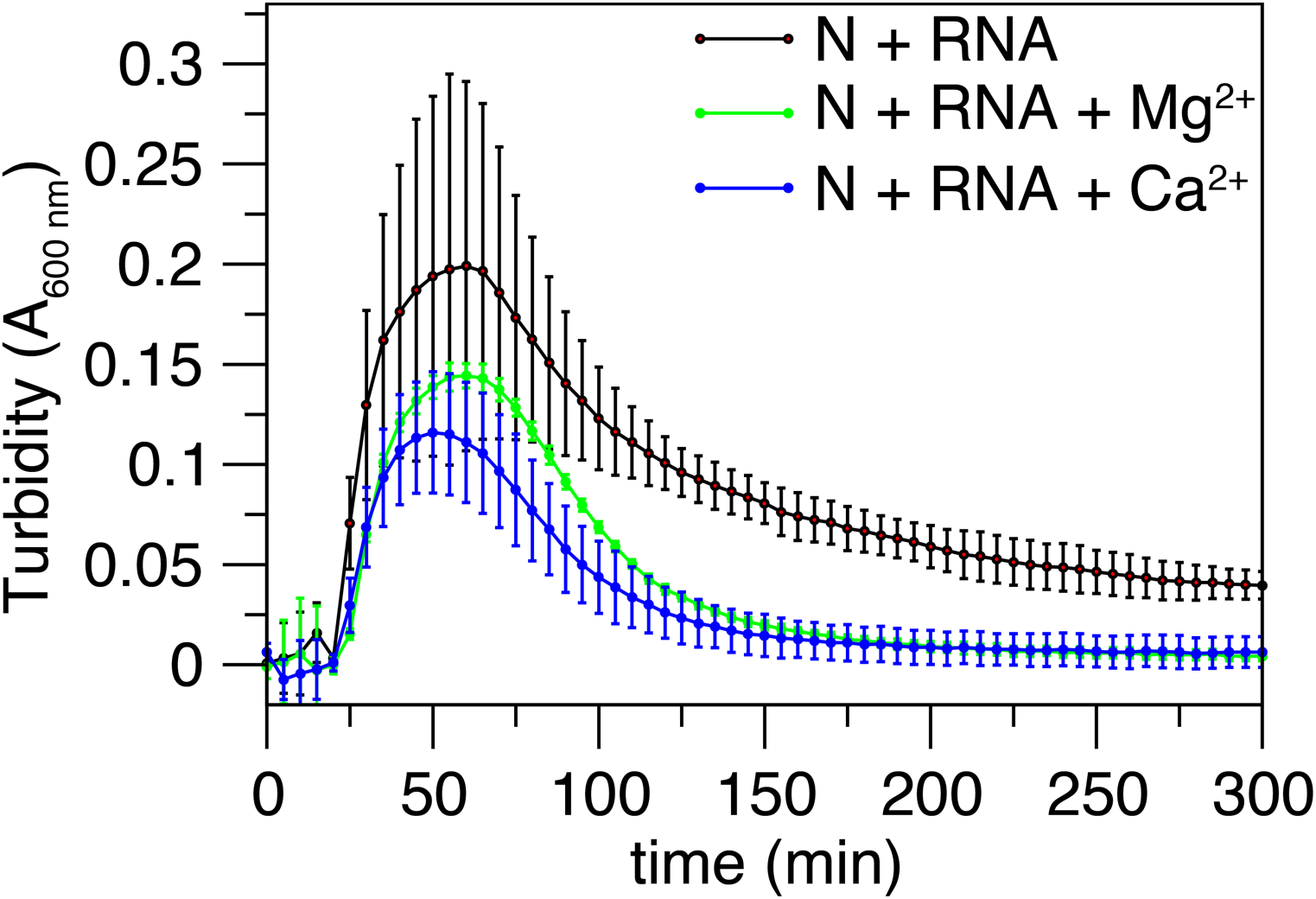
Divalent metal salts do not substantially alter N LLPS. Addition of 2 mM MgCl_2_ or CaCl_2_ does not alter LLPS of 50 μM MBP-N in the presence of 0.5 mg/mL RNA and 70 mM NaCl.

